# The dichotomous impacts of semaphorin3E deficiency on exacerbating airway hyperresponsiveness, remodelling, and inflammation in type-2 low and type-2 high asthma models

**DOI:** 10.1101/2025.03.23.644815

**Authors:** Mojdeh Matloubi, Fatemeh Sedaghat, Lianyu Shan, Sujata Basu, Andrew Halayko, Abdelilah S. Gounni

## Abstract

Semaphorin3E has shown promise in alleviating the severity of asthma in preclinical studies; however, its role in the chronic features of type 2-low asthma remains unclear. Therefore, we aimed to investigate the role of Sema3E in a mouse model of severe asthma that exhibits a mix of granulocytic inflammation with neutrophils dominance and compared the results with those from the type-2 high eosinophilic asthma model.

*Sema3E knockout (KO)* and wild-type (WT) mice were subjected to type-2 low and type-2 high regimens using house dust mite (HDM) combined with cyclic-di-GMP or HDM alone, respectively. Airway hyperresponsiveness parameters were measured using the FlexiVent ventilator. Bronchoalveolar lavage fluid cell phenotyping was performed by flowcytometry. Additionally, cytokines and antibodies were quantified using Mesoscale and ELISA. Mucus overproduction and goblet cell hyperplasia were visualized by Periodic-acid–Schiff staining.

In comparison to WT mice, *Sema3E KO* mice exhibited an enhanced tissue resistance and tissue elastance in the type 2-low asthma model. Concurrently, *Sema3E KO* mice that were subjected to the type-2 low asthma model demonstrated an elevated presence of pulmonary neutrophils, dendritic cells, CD4 T cells, as well as increased levels of IL-17, TNF, IL-1β, CXCL-8, and MCP-1/CCL2 in comparison to their WT counterparts. However, in the type-2 high model, *Sema3E KO* mice exhibited a significant increase in goblet cell numbers and mucus overproduction, as well as enhancements in the number of eosinophils, IgE-producing B cells, and IL-4 levels compared to WT mice, highlighting the homeostatic role of Sema3E in the distinct immune niche of type-2 low and type-2 high asthma. Overall, our data showed that Sema3E is critical in modulating AHR, airway inflammation, and tissue remodelling in type 2 low and type 2 high phenotypes of asthma. The Sema3E regulatory network varies depending on the immunization regimen, affecting distinct parameters in type-2 low and type-2 high asthma models.

## Introduction

Asthma is a complex condition with various observable features, known as phenotypes, that arise from specific biological mechanisms called endotypes (1, 2). Each phenotype exhibits its own severity, history, and medication response, influenced by associated biological mechanisms (1, 2). Type-2 low (neutrophilic) and type-2 high (eosinophilic) asthma represent two distinct inflammatory phenotypes within the broader spectrum of asthma (3–5). Neutrophilic asthma (type 2 low) is characterized by a predominant presence of neutrophils in the airways and an immune response marked by elevated levels of inflammatory cytokines, such as IL-17 and IFN-γ, manifesting as a more severe and persistent form of the disease (3–5). Conversely, eosinophilic asthma, classified as type-2 high asthma, is commonly associated with allergic triggers and is characterized by a significant presence of eosinophils within the airways, and associated with elevated levels of cytokines, like IL-4, IL-5, IL-13, and IgE (3–5). Treatment strategies may be tailored based on the predominant inflammatory cell type, with corticosteroids often effective in addressing eosinophilic inflammation (3–5). The distinction between these phenotypes is crucial for personalized asthma management, considering the diverse underlying mechanisms contributing to airway inflammation and remodelling in different individuals.

Semaphorins were initially identified as axon guidance during the development of the neural system (6). However, they are present in different organs and tissues, participating in diverse signaling pathways (7, 8). In respiratory diseases, semaphorins play crucial roles in cell-cell contact, differentiation, proliferation, migration, and immune system regulation (9–13).

Our lab demonstrates that Sema3E plays a modulatory role in allergic HDM model of asthma, and the Sema3E-plexinD1 signaling to be critical for maintaining homeostatic conditions within the airways (14–26). The deficiency of Sema3E in murine models of allergic asthma resulted in a significant infiltration of granulocytes into the pulmonary tissue, along with an amplified Th2/Th17 immune response. This condition was characterized by pronounced airway hyperresponsiveness (AHR), excessive mucus and collagen production, as well as hyperplasia and hypertrophy of smooth muscle cells (14, 22, 23, 26). In contrast, the intranasal administration of exogenous Sema3E reduced these pathological effects (14, 22, 23, 26), hence underscoring the essential role of the Sema3E-plexinD1 axis in maintaining homeostasis in allergic asthma (14). However, the role of Sema3E in type 2 low asthma has not been investigated.

In this regard, our lab has established two chronic allergen protocols that induce robust airway granulocytic inflammation associated with airway hyperresponsiveness (AHR) and remodelling (27). Given the potential of Sema3E as a therapeutic target for allergic HDM asthma (14, 15), we sought to investigate the role of Sema3E in type 2-low asthma using preclinical models and to compare the effects of Sema3E deficiency in this model with those observed in the type 2-high asthma phenotype.

Our findings reveal that Sema3E plays a pivotal role in regulating airway hyperresponsiveness (AHR), inflammation, and tissue remodeling across different asthma phenotypes. In the type 2-low asthma model, Sema3E-deficient mice exhibited increased tissue resistance and elastance, accompanied by elevated levels of neutrophils, dendritic cells, CD4+ T cells, and pro-inflammatory cytokines such as IL-17, TNF, IL-1β, CXCL-8, and MCP-1/CCL2. In contrast, Sema3E deficiency in the type 2-high model led to a marked increase in goblet cell hyperplasia, mucus overproduction, eosinophilia, B cell producing IgE, and elevated IL-4 levels. These results suggest that Sema3E contributes to maintaining homeostasis within distinct immune niches, exerting differential effects in neutrophil-dominant versus eosinophil-dominant asthma. Overall, our study reveals that the Sema3E regulatory network is highly context-dependent, underscoring its pivotal role in orchestrating immune and structural responses across distinct asthma phenotypes. These findings highlight Sema3E as a key immunoregulatory molecule with potential implications for phenotype-specific therapeutic targeting in asthma.

## Method and materials

### Animals

Eight-week-old male and female Balb/c *Sema3E knocked out (KO)* mice were generated by backcrossing *Sema3E KO* (129 P2) for ten generations (generously provided by Dr. Fanny Mann, Developmental Biology Institute of Marseille Luminy, Université de la Méditerranée, Marseille, France). The littermate wild-type Balb/c mice were used as control (9).

All mice were kept in the pathogen-free room at the Central Animal Care facility, University of Manitoba. All procedures followed the guidelines provided by the Canadian Council for Animal Care and approved by the University of Manitoba Animal Care and Use Committee (protocol number 19-035).

### Allergen-induced airway inflammation model

Eight-week-old male and female wild-type and *Sema3E KO* mice underwent chronic neutrophilic (type-2 low) and eosinophilic (type-2 high) asthma regimens for four weeks (Fig 1A). For the severe type-2 low model, mice underwent sensitization with 25 μg of low-endotoxin house dust mite (HDM) (lot 259585; Der-p-1: 298.36 mcg/vial, protein: 5.34 mg/vial, endotoxin: 615 EU/vial, Greer Laboratories, Lenoir, NC) and 5 μg of cyclic di-GMP (Cat# SML1228-1UMO; Millipore Sigma) intranasally on days 1, 3, and 5. Following a five-day resting period, the mice were subjected to three challenge sets, each comprising three consecutive challenges with HDM and c-di-GMP, with a four-day rest interval between each set. On the first day of each challenge set, 0.5 μg of c-di-GMP was administered in conjunction with HDM, followed by two subsequent challenges that consisted solely of 25 μg of HDM (Fig 1A) (27). The eosinophilic model followed the same sensitization and challenge protocol, except that c-di-GMP was not used in this model (27). The mice were sacrificed 24 hours after the final challenge to investigate the outcomes. *Sema3E KO* and wild-type control mice, which did not receive allergens, were used as control mice in this study.

**Fig 1.**
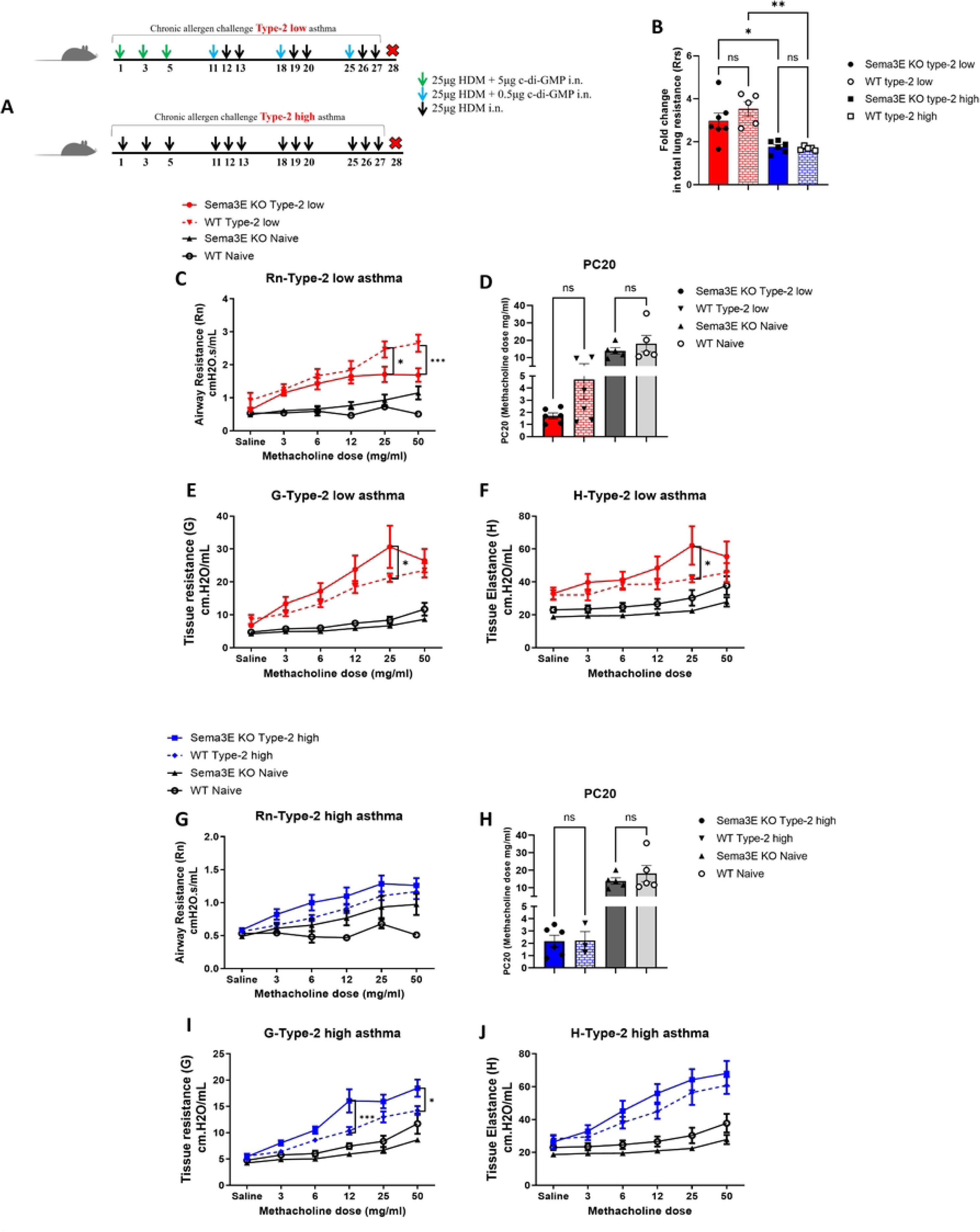
Sema3E ablation highly enhanced bronchoconstriction in type-2 low compared to type-2 high model of asthma. **(A)** The type-2 low and type-2 high models of asthma were established by intranasal exposure to c-di-GMP+HDM or HDM alone for four weeks, respectively. *Sema3E KO* and WT mice underwent tracheotomy and were subsequently received increasing gradients of methacholine to measure airway hyperresponsiveness parameters. **(B)** Fold increase in total lung resistance (Rrs) was calculated for each *Sema3E KO* and WT group (type-2 low and type-2 high) relative to the mean of their respective naïve counterparts. **(C & G)** airway resistance, **(D & H)** PC20, **(E & I)** tissue resistance, and **(F & J)** tissue elastance in both type −2 low and type-2 high models of asthma were calculated between *Sema3E KO* and WT mice. All data shown are representative of 3-7 mice per group. Data are presented as the mean with standard error of the mean (SEM), 2-way ANOVA *p < 0.05, ***p < 0.001

### Methacholine challenge test

Airway hyperresponsiveness (AHR) was assessed by measuring airway resistance (Rn), tissue resistance (G), and tissue elastance (H) using the FlexiVent animal ventilator (Scireq, Montreal, QC, Canada). Both allergen-exposed and control mice underwent thoracotomy, followed by intratracheal administration of methacholine at progressively increasing concentrations (Saline, 3, 6, 12, 25, and 50 mg/ml) in 5-minute intervals. Lung function assessments were conducted as previously described (22).

### Bronchoalveolar lavage fluid collection and differential cell count

Bronchoalveolar lavage fluid (BALF) was obtained from the airways through two administrations of 1 ml sterile PBS containing 0.05 mM EDTA. After centrifugation, the supernatant was stored at −80◦C for subsequent analysis. The total number of cells in BALF was counted using trypan blue and a hemocytometer. Subsequently, cells were processed through cytospins, fixed, and stained with Wright-Giemsa (22).

### Immunophenotyping of lung and BALF immune cells

Lung tissue from *Sema3E KO* and WT mice were collected, and single-cell suspensions were prepared as previously described (26). Following washing and incubation with Fc-blocker, cells were stained with a mixture (0.5μl of antibodies/20μl of flow buffer per tube) containing the following anti-mouse antibodies using four antibody panels. The first panel interrogating granulocytes composition, consisted of fixable viability dye eFluor 780 (eBioscience), CD45-PB (clone S18009F; BioLegend), MerTK-APC (clone 2B10C42; BioLegend), CD64-PerCP/Cy5.5 (clone X54-5/7.1; BioLegend), CD11b-PE/Cy7 (clone M1/70; eBioscience), CD11c-APC/Cy7 (clone N418; BioLegend), Siglec F-PE (clone E50-2440; BD Biosciences), Gr-1-FITC (clone RB6-8C5; BioLegend). The second panel, used to probe for lung myeloid conventional dendritic cells, consisted of dumping antibodies, including CD3-PE (clone 17A2; BioLegend), B220-PE (clone RA3-6B2; BioLegend), NKp46-PE (clone 29A1.4; BioLegend), Gr-1-PE (clone RB6-8C5; BioLegend), and Siglec-F-PE (clone E50-2440; BD Biosciences) to remove lymphocytes, NK, and granulocytes. Followed by fixable viability dye eFluor 780 (eBioscience), CD45-APC-Cy7 (clone 30-F11; BioLegend), MHC class II (I-A/I-E)-PB (clone AF6-120.1; BioLegend), CD11c-FITC (clone N418; BioLegend), CD103-PerCP/Cy5.5 (clone 2E7; BioLegend), CD11b-PE-Cy7 (clone M1/70; eBioscience).

The third panel, is used to analyze lung B cell and intracellular staining for antibodies, consist of fixable viability dye eFluor 780 (eBioscience), CD45-PB (clone S18009F; BioLegend), CD19-APC-Cy7 (clone 1D3/CD19; BioLegend), B220-APC (clone RA3-6B2; BioLegend), IgE-FITC (clone RME-1; BioLegend), IgG1-PE-Cy7 (clone RMG1-1; BioLegend),).

The fourth panel, CD4+ and CD8+ T cells, consists of fixable viability dye eFluor 780 (eBioscience), CD45-APC-Cy7 (clone 30-F11; BioLegend), CD3-PE-Cy7 (clone 17A2; BioLegend), CD4-PB (clone GK1.5; BioLegend), CD8-FITC (clone 53-5.8; BioLegend).

Moreover, inflammatory cells in the BALF were characterized using anti-mouse antibodies, including MerTK-APC (clone 2B10C42; BioLegend), CD64-PerCP/Cy5.5 (clone X54-5/7.1; BioLegend), CD11b-PE/Cy7 (clone M1/70; eBioscience), CD11c-APC/Cy7 (clone N418; BioLegend), Siglec F-PE (clone E50-2440; BD Biosciences), Gr-1-BV421 (clone RB6-8C5; BioLegend), CD4-BV605 (clone GK1.5; BioLegend), and B220-BV500 (clone RA3-6B2; eBioscience). Subsequently, the samples were acquired using the Beckman Coulter CytoFLEX flow cytometer and analyzed using FlowJo software.

### Cytokine measurement

The level of TNF, IL-1β, IL-4, IL-5, IL-10, IL-13, IL-17A, CXCL-8 and IFN-γ was measured using MesoScale Discovery system (MSD U-PLEX, US. Cat#K15069M-1) in BALF supernatants according to the manufacturer’s instructions. Plates were read by MESO QuickPlex SQ 120MM machine (Rockville, Maryland, USA).

### Measurement of Immunoglobulins in serum

Cardiac blood was collected from Sema3E KO and wild-type mice exposed to either saline or HDM. After centrifugation, serum samples were isolated to quantify total and HDM-specific immunoglobulins using ELISA, following the manufacturer’s protocols (22, 28). Antibodies for total Ig detection were obtained from Southern Biotech (Birmingham, AL). ELISA data were analyzed using SoftMax Pro software (Molecular Devices).

### Lung Histology

The left lung lobe was collected and fixed in 10% formalin for 24 hours, then transferred to 70% alcohol. Next, the lung tissue was processed by gradually increasing the alcohol concentration from 70% to 100%, followed by two xylene washes, and then embedded in paraffin. The tissue was sectioned at 5 µm using a microtome. The quantification of mucus production in lung tissue sections was investigated by performing periodic acid Schiff (PAS), (22, 29) and the results were quantified using ImageJ software. Briefly, the length of the basal membrane was determined by assigning the region of interest with the freehand tool. Using the color difference between the PAS-stained goblet cells (purple) and other basal membrane cells, the software counted the number of goblet cells within the specified region of interest. Moreover, inflammation severity in the lungs was assessed using H&E staining, and the results were reported as a pathological score, as previously described. (30).

### Statistical analysis

GraphPad Prism 9.0 software was used for statistical analysis. Depending on the number of groups and treatments, data were analyzed by two-way ANOVA, followed by a Tukey test. Differences were statistically significant at *p < 0.05, **p < 0.01, and ***p < 0.001, ****p < 0.0001.

## Results

### Sema3E deficiency exaggerated bronchoconstriction in type-2 low model of asthma

The key clinical hallmark of asthma is airway hyperresponsiveness (AHR), which arises from abnormalities in the structural cells of the lungs along with enhanced airway inflammation (31). Initially, we assessed the impact of Sema3E deficiency on baseline and allergen-induced AHR in type-2 low and compared to type-2 high models of asthma through the evaluation of lung function parameters using the methacholine test (22).

*Sema3E KO* and wild-type mice were exposed to c-di-GMP+HDM or HDM alone for four weeks, as outlined in the model plan (Fig 1A), to induce mixed granulocytic with neutrophil dominance and eosinophil dominance asthma models, respectively. Both the c-di-GMP+HDM and HDM alone challenge significantly increased the constrictive response of the airways to all doses of methacholine. As shown in Fig. 1B, total lung resistance (Rrs), a parameter reflecting overall lung function, is increased three-fold in *Sema3E KO* and WT type-2 low mice compared to naïve counterparts. Additionally, Rrs is elevated two-fold in both *Sema3E KO* and WT type-2 high mice relative to naïve controls, indicating that type-2 low asthma presents a more severe phenotype in terms of physiology and lung function than the type-2 high model (p<0.05) (Fig. 1B).

Airway resistance is one of the parameters of AHR defining as the resistance encountered by airflow in the airways, especially the smaller bronchioles (32). Enhanced airway resistance suggests narrowed air passages, which can stem from inflammation, constriction of smooth muscles, and increased mucus production, ultimately causing breathing challenges (32). Our results revealed that airway resistance (Rn) did not increase in response to Sema3E deletion compared to WT mice in the type-2 low asthma model (Fig. 1C). Further, there was also a tendency to higher constrictive responses in the WT type-2 low model compared to *Sema3E KO* mice, although this difference did not reach statistical significance (Fig 1C). This phenomenon may indicate that, due to the severity of the disease in our model, the airways no longer respond to methacholine. This could be interpreted as increased hypersensitivity of the airways to the methacholine dose, rather than hyperreactivity (33) in type-2 low asthma following Sema3E deletion. Therefore, we investigated the PC20 (provocative concentration of a substance that causes a 20% decrease in a specific physiological parameter). A 20% fall in FEV1 in response to methacholine challenge (PC20) is equivalent to a 43.5% rise in respiratory resistance (34). Thus, in our model, the methacholine PC20 was calculated by assessing the methacholine concentration required to increase airway resistance by 43.5% from the baseline. As shown in Figure 1D, *Sema3E KO* mice in the type 2-low asthma model exhibited an increasing trend to sensitivity to methacholine. The amount of methacholine required to induce 43.2% airway resistance (Rn) was lower in *Sema3E KO* mice (PC20 ≈ 2 mg/ml) compared to their wild-type counterparts (PC20 ≈ 4 mg/ml) and was reduced eightfold compared to naïve *Sema3E KO* and WT control mice (PC20 ≈ 16 mg/ml). These findings indicate that *Sema3E KO* mice develop heightened airway sensitivity to methacholine, rather than increased airway reactivity, in the type 2-low asthma model.

Other parameters of AHR, such as tissue resistance (G) and tissue elastance (H) were investigated to better understand the lung function in *Sema3E KO* and WT mice in both type-2 low and type-2 high asthma models. Both tissue resistance (G) and tissue elastance (H) showed a statistically significant increase in the *Sema3E KO* type-2 low asthma model that peaked at the 25 mg/ml methacholine dose (p<0.05) compared with the WT counterpart (Fig 1E, 1F).

In the type-2 high asthma model, although *Sema3E KO* mice did not display statistically significance exaggerated airway resistance (Rn) compared to WT mice, there is a higher trend in airway resistance and hyperreactivity in *Sema3E KO* compared to WT counterpart (Fig 1G). Moreover, PC20 highlights the same hyperreactivity of airways to methacholine dose between *Sema3E KO* and WT in type-2 high (PC20 ≈2 mg/ml) compared to naïve control groups (PC20≈16 mg/ml) (Figure 1H).

Furthermore, tissue resistance (G) demonstrated a significant increase in the *Sema3E KO* type-2 high asthma model that peaked at the 12 mg/ml (p<0.001) and dose 50 mg/ml methacholine dose (p<0.05) compared with the WT counterpart (Figure 1I). In response to Sema3E ablation, tissue elastance (H) showed a higher tendency in all doses of methacholine compared to WT mice, despite that it did not reach the statistical significance (Figure 1J). However, the Rrs parameter, which represents total lung function, was significantly increased in response to Sema3E deletion in the type 2-high model. This finding highlights that lung function was significantly compromised in *Sema3E KO* mice compared to their WT counterparts in the type 2-high model (data not shown).

Overall, these findings demonstrate that the global ablation of Sema3E led to more exaggerated bronchoconstriction and poor lung function in type-2 low compared to type-2 high asthma model. This is evidenced by increased airway hyperresponsiveness and hypersensitivity of the lungs to methacholine.

### Sema3E deficiency selectively enhanced inflammatory cell recruitment into the lungs of both type-2 low and type-2 high asthma

Inflammation plays a central role in the pathophysiology of asthma, contributing to the chronic nature of this respiratory condition (35). We assessed the effect of Sema3E deletion on immune cell composition or recruitment into the lungs using flow cytometry in both type-2 low and type-2 high models of asthma.

The influx of inflammatory cells in the lungs was markedly induced by allergen challenge, reaching 0.8-1.1 x 10^6^ cells/ml in the BALF in the type-2 low model and 0.3-0.5 x 10^6^ cells/ml in type-2 high asthma model compared to the and naïve mice (0.03-0.06 x 10^6^ cells/mL) (Fig 2C). Also, our fold change analysis confirmed a five-fold increase in the total number of immune cells in *Sema3E KO* type-2 low mice compared to their naïve counterparts. Similarly, we observed a three/four-fold increase in the WT type-2 low model compared to its naïve counterpart, indicating that Sema3E deletion exacerbates lung inflammation (p<0.05) (Fig. 2D). Moreover, both the *Sema3E KO* and WT type-2 high models showed a two-fold increase in total immune cells compared to their naïve counterparts. These findings suggest that the type-2 low model exhibits a more severe phenotype in terms of lung inflammation compared to the type-2 high model (Fig 2D).

**Fig 2.**
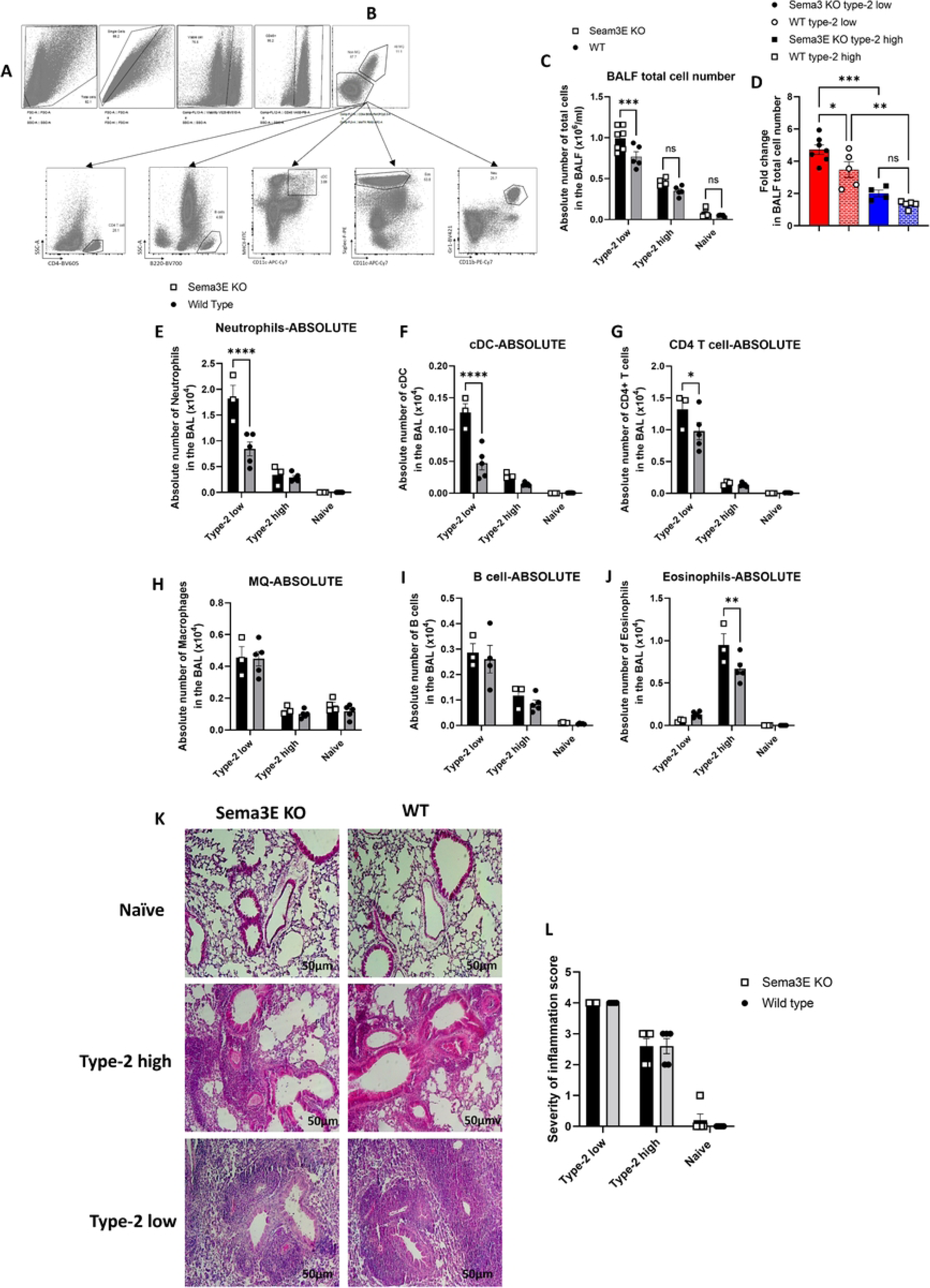
Sema3E ablation on chronic allergic airway inflammation differed based on the immunization regimen. Bronchoalveolar lavage fluid (BALF) was collected from Sema3E KO and wild-type mice to analyze immune cell populations using flow cytometry (FACS) in both type-2 low and type-2 high asthma models. **(A)** The gating strategy involved removing debris and doublets, followed by selecting viable leukocytes (CD45+). **(B)** Macrophages were identified by co-expression of MerTK and CD64, while non-macrophage populations (lacking MerTK and CD64) were further analyzed to distinguish other inflammatory cell subsets. Neutrophils were characterized by Ly6G+ (1A8) and CD11b+ expression, whereas eosinophils were defined as Siglec-F+/CD11c-. Conventional dendritic cells (cDCs) were identified based on MHCII and CD11c co-expression within the non-macrophage population. Additionally, CD4 T cells and B cells were characterized by surface expression of CD4 and B220, respectively. **(C)** The total cell count, **(D)** fold change, and differential immune cell populations, including **(E)** neutrophils, **(F)** cDCs, **(G)** CD4 T cells, **(H)** macrophages, **(I)** B cells, and **(J)** eosinophils, were compared between Sema3E KO and WT mice in both asthma models. **(K)** Inflammation severity was assessed through H&E staining of paraffin-embedded lung sections, and **(L)** pathological scores were reported. Scale bars represent 50 µm. Data are representative of 3–7 mice per group and are presented as mean values (pre-gated on CD45+) with standard error of the mean (SEM). Statistical significance was determined using two-way ANOVA (**p < 0.01, ***p < 0.001, ****p < 0.0001).

In the type-2 low asthma model, we observed more immune cell recruitment and severe inflammatory responses with a dominance of neutrophils (Fig 2C, 2D, and 2E). However, in the type-2 high model, we observed milder responses associated with eosinophils (Fig 2C, 2D, 2J). These findings highlight the neutrophilic endotype of type-2 low asthma model and eosinophilic endotype of type-2 high model of asthma.

In the type-2 low asthma model, we observed a higher number of neutrophils (p<0.0001), dendritic cells (DC) (p<0.0001), and CD4+ T cells (p<0.05) in *Sema3E KO* mice compared to WT mice (Fig 2E, 2F, 2G). However, the number of other inflammatory cells, including macrophages, B cells, and eosinophils, was not significantly different between *Sema3E KO* and WT mice in the type-2 low model (Fig 2H, 2I, 2J).

In type-2 high asthma, the number of inflammatory cells, including neutrophils, cDC, CD4 T cells, macrophages, and B cells did not change in *Sema3E KO* mice compared to WT counterparts. However, eosinophils were the cell type that significantly accumulated in the lungs of *Sema3E KO* mice compared to WT mice in the type-2 high model (p<0.001) (Fig 2J).

Moreover, H&E staining of lung tissue sections demonstrated more severe inflammation in type-2 low model compared to type-2 high. However, we did not observe any significant changes in severity of inflammation in *Sema3E KO* mice compared to their WT counterparts in both type-2 low and type-2 high models of asthma (Fig. 2K & 2L).

In summary, in the type-2 low model, we observed more immune cell recruitment and severe responses with a dominance of neutrophils, CD4+ T cells and DC in the absence of Seam3E. However, in the type-2 high model, we observed milder responses associated with eosinophilia. Overall, the global ablation of Sema3E enhanced cell migration into the lungs based on each allergen regimen. However, this deletion did not affect the number of macrophages and B cells in either of the asthma models.

### Sema3E ablation selectively increased cytokine production in asthma based on the allergen regimen

Both type-1 and type-2 inflammatory cytokines play a critical role in asthma pathogenesis and its progression (36). Accordingly, we investigated the levels of pro-/anti-inflammatory cytokines in *Sema3E KO* and WT counterparts in both type-2 low and type-2 high models of asthma.

Compared to naïve mice, allergen challenge induced a significant increase in inflammatory cytokines and chemokines levels in both type-2 low and type-2 high asthma models. We could not detect IFN-γ, IL-17, IL-4, IL-5, IL-10, and MCP-1/CCL2 in naïve mice, but after the allergen challenge, their levels enhanced significantly in both models selectively (Fig. 3). As expected, the levels of type-1 cytokines, such as IFN-γ (40-80 pg/ml) and IL-17 (20-40 pg/ml), were higher in the type-2 low model of asthma compared to the type-2 high model (0.1-0.4 pg/ml and 1-4 pg/ml, respectively) (Fig. 3A, 3B). However, the levels of type-2 cytokines, such as IL-4 (10-25 pg/ml), IL-5 (10-16 pg/ml), and IL-13 (30-70 pg/ml), were enhanced in response to the type-2 high model compared to the type-2 low asthma model (0.5-3 pg/ml, 1-4 pg/ml, and 2-8 pg/ml, respectively) (Fig. 3C, 3D, 3E).

**Fig 3.**
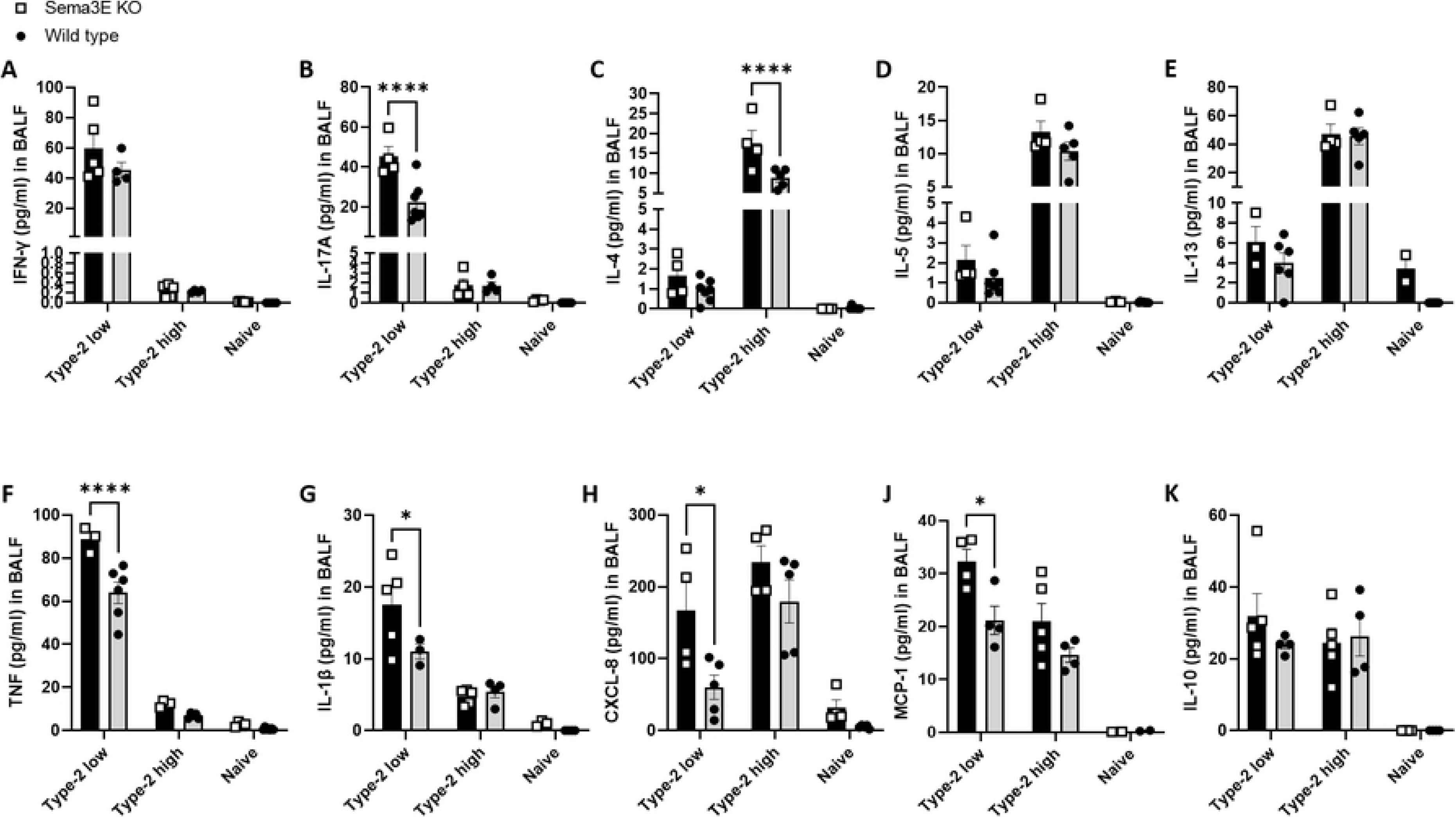
Sema3E deletion enhanced the levels of inflammatory cytokines in type-2 low and type-2 high asthma models. The levels of **(A)** IFN-γ, **(B)** IL-17A, **(C)** IL-4, **(D)** IL-5, **(E)** IL-13, **(F)** TNF, **(G)** IL-1β, **(H)** CXCL-8, **(J)** MCP-1/CCL2, and **(K)** IL-10 were measured using mesoscale in BALF supernatants obtained from *Sema3E KO* and WT mice after intranasal exposure to allergen in both type-2 low and type-2 high models of asthma. All data shown are representative of 3-7 mice per group. Data are presented as the mean with the standard error of the mean (SEM), 2-way ANOVA, *p < 0.05, ****p < 0.0001.

We observed higher levels of IL-17 (p<0.0001), TNF (p<0.0001), IL-1β (p<0.05), CXCL-8 (p<0.05), and MCP-1/CCL2 (p<0.05) in *Sema3E KO* mice compared to WT mice in the type-2 low model (Fig. 3B, 3F, 3G, 3H, 3J). However, the levels of IFN-γ, IL-10, and type-2 cytokines, including IL-4, IL-5, and IL-13, were not significantly different between *Sema3E KO* and WT mice in the type-2 low asthma model (Fig. 3A, 3C, 3D, 3E, 3K). On the other hand, in the type-2 high model, the level of IL-4 (p<0.0001) was significantly higher in *Sema3E KO* mice compared to WT mice (Fig. 3C), although there was no significant difference in the levels of IFN-γ, IL-17, IL-5, IL-13, TNF, IL-1β, CXCL-8, MCP-1/CCL2, and IL-10 between *Sema3E KO* and WT mice in the type-2 high model of asthma (Fig. 3A-3K).

In summary, Sema3E deficiency in type-2 low asthma mediates more severe airway inflammation and exaggerated type-1 cytokines profile, while in the type-2 high it induced enhanced type-2 cytokines and milder airway inflammation. Hence, our data suggests that Sema3E selectively regulates cellular and molecular niches in both chronic asthma models.

### Sema3E ablation had divergent effects on antibody-producing B cell responses depending on asthma subtypes

We measured the levels of total and HDM specific antibodies and the number of B cells producing antibodies in *Sema3E KO* and WT counterparts in both type-2 low and type-2 high models of asthma.

In naïve mice, the levels of total IgG1 (14-80 µg/ml) and total IgE (300-800 ng/ml) were very low. However, in response to allergen challenge, the total IgG1 levels increased to 800-1200 µg/ml in the type-2 low model and 120-300 µg/ml in the type-2 high model of asthma. Moreover, the levels of total IgE increased to 5000-10,000 ng/ml in type-2 low model and 15,000-20,000 ng/ml in type-2 high model of asthma (Fig. 4A & 4C). Furthermore, we found higher levels of HDM-specific IgG1 in type-2 low model, while the levels of HDM-specific IgE were higher in the type-2 high model (Fig. 4B & 4D), highlighting the different immune niches and responses between these two models. However, in response to Sema3E deletion, the total and HDM specific serum IgG1 and IgE levels did not change compared to WT mice in both models of asthma (Fig. 4A-4D).

**Fig 4.**
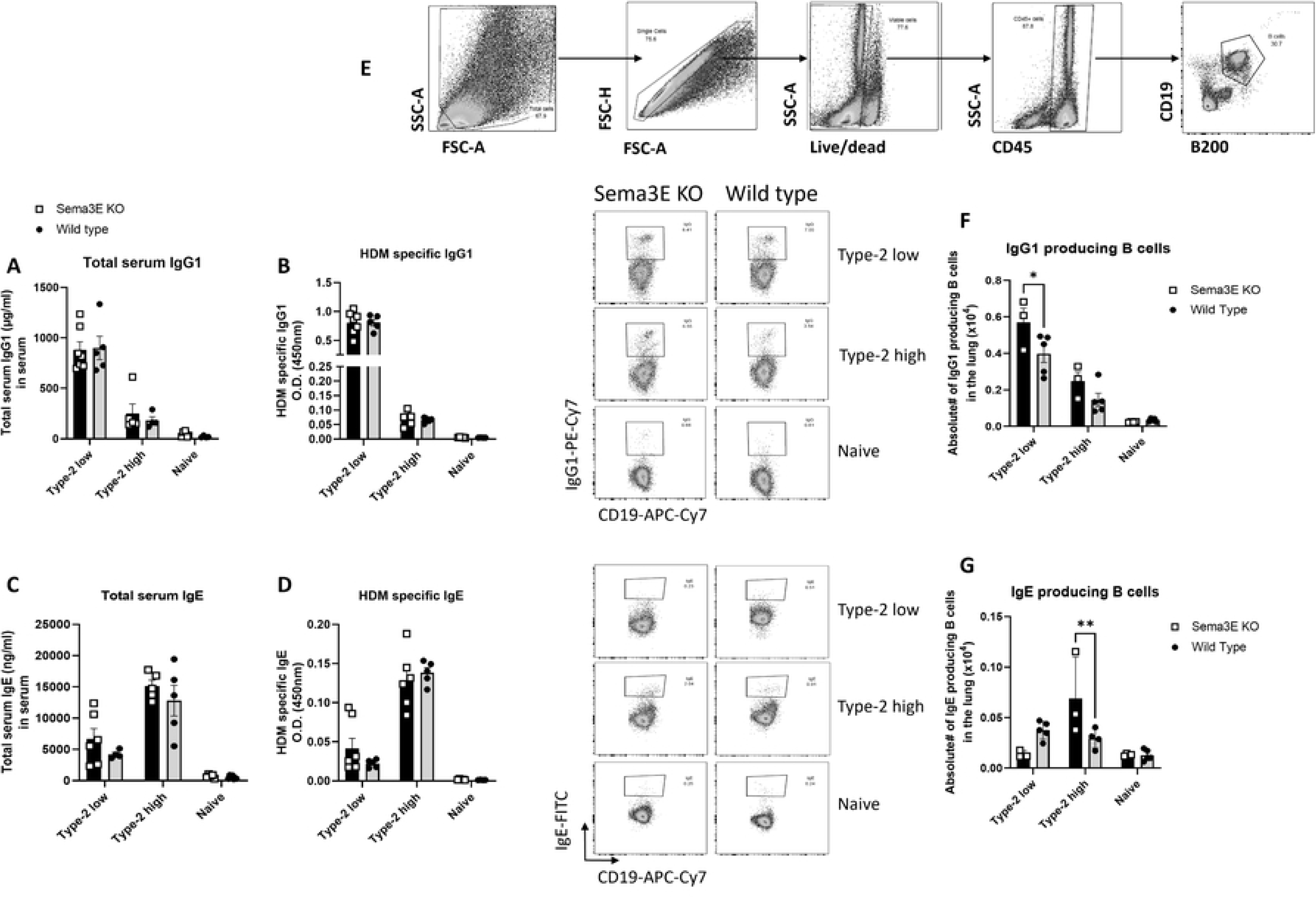
Sema3E deficiency has differential effects on B cell responses in distinct asthma phenotypes. Serum samples were obtained from Sema3E KO and WT mice following allergen immunization in type-2 low and type-2 high asthma models. The concentrations of **(A & B)** total and HDM-specific IgG1 and **(C & D)** IgE were quantified using ELISA. Additionally, flow cytometry (FACS) was performed on lung single-cell suspensions from Sema3E KO and WT mice to analyze B cell-derived antibody production. **(E)** The gating strategy involved selecting total and single cells, followed by identifying viable CD19+ and B220+ double-positive cells as B lymphocytes within the CD45+ population. Antibody production by B cells was assessed by gating on CD19+ cells, and the levels of **(F)** IgG1 and **(G)** IgE were measured. All data represent findings from 3– 7 mice per group and are presented as mean values (pre-gated on CD45+) with standard error of the mean (SEM). Statistical significance was determined using two-way ANOVA (*p < 0.05, **p < 0.01).

Furthermore, we investigated the number of B cells producing antibodies in the lungs of both type-2 low and type-2 high models of asthma using flow cytometry. We showed that the number of IgG1-producing B cells (p<0.05) significantly increased in *Sema3E KO* mice compared to WT mice in the type-2 low model (Fig. 4D). Moreover, the number of IgE-producing B cells increased in the type-2 high model (p<0.001), specifically in *Sema3E KO* mice compared to WT mice (Fig. 4E).

Overall, the levels of IgG and IgE increased selectively in response to Sema3E deficiency, reflecting the unique immune niche in each chronic asthma model and emphasizing the homeostatic role of Sema3E in these contexts.

### Sema3E deletion significantly increased the number of goblet cells and mucus production in type-2 high asthma model

Mucus overproduction, one of the hallmarks of asthma, leads to airway thickening and permanent narrowing, which in turn reduces lung function by increasing airflow resistance, impairing gas exchange, and exacerbating respiratory symptoms (37). We investigated goblet cell hyperplasia and mucus production using Periodic acid-Schiff (PAS) staining (Fig 5A) in *Sema3E KO* and WT counterparts in both type-2 low and type-2 high models of asthma.

**Fig 5.**
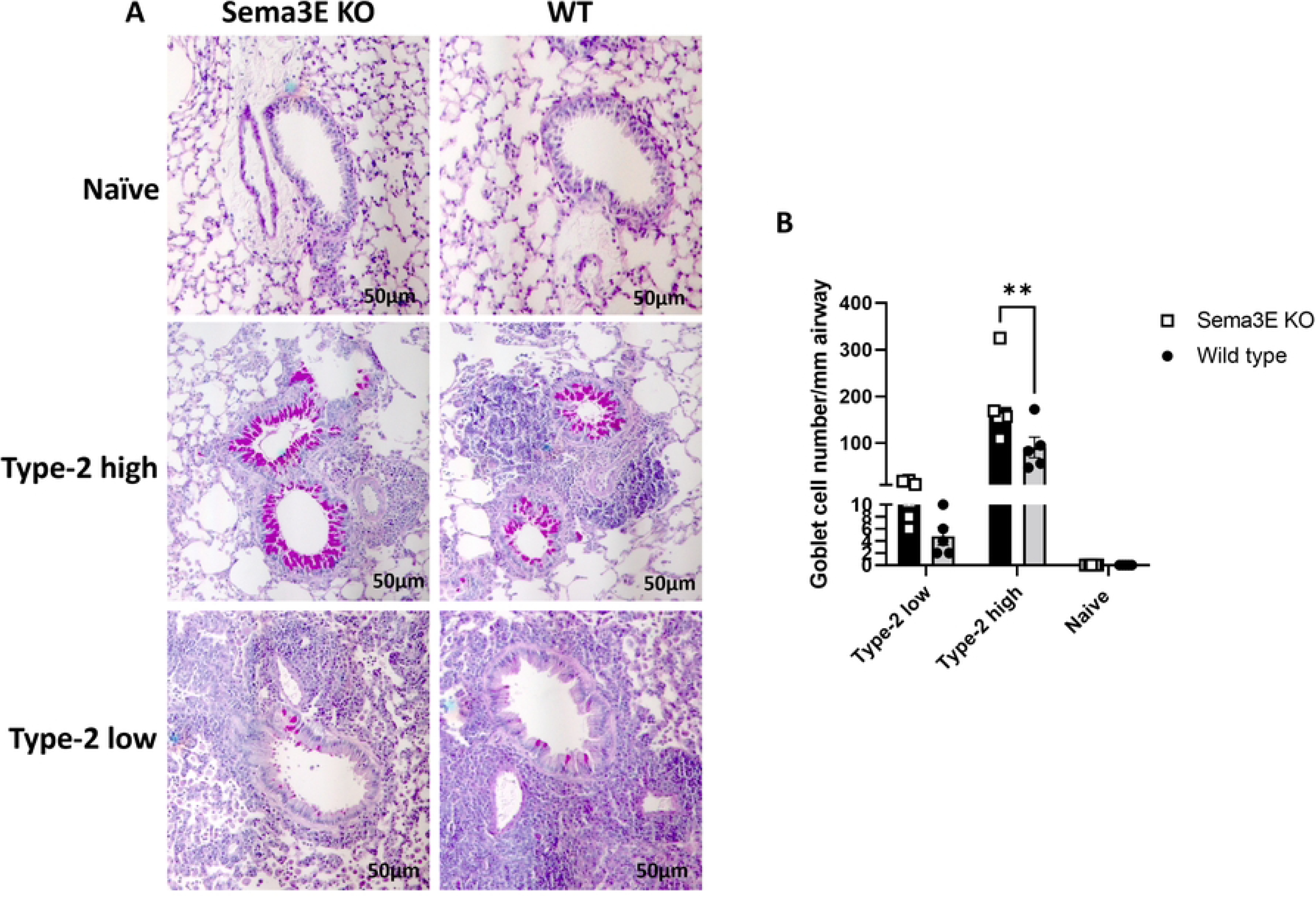
Sema3E deficiency increased goblet cells hyperplasia and mucus overproduction in type-2 high model of asthma. **(A)** Goblet cell hyperplasia/mucus production and was visualized using periodic acid-Schiff (PAS) staining in the paraffin-embedded lung sections. **(B)** The result were analyzed by imageJ software. Scale bars, 50µm; All data shown are representative of 3-7 mice per group; 2-Way ANOVA, **p<0.01.

In naïve mice, the amount of mucus was very low and undetectable in both *Sema3E KO* and WT mice. However, allergen challenge induced goblet cell hyperplasia and mucus production, with 3-10 goblet cells/mm of the basement membrane detected in the type-2 low asthma model and 100-200 goblet cells/mm of the basement membrane in the type-2 high asthma model (Fig 5B).

Sema3E ablation significantly increased goblet cell hyperplasia compared to WT mice (p<0.01) in the type-2 high asthma model (Fig 5A). Although a similar trend was observed in the type-2 low model, the results did not reach statistical significance (Fig 5A).

Overall, Sema3E ablation significantly increases goblet cell hyperplasia and mucus production in the type-2 high asthma model. However, in the type-2 low model, mucus production did not show a statistically significant increase. This suggests that Sema3E plays a regulatory role in mucus production, highlighting its selective action across different tissue niches and asthma phenotypes.

## Discussion

Asthma mouse models are designed to replicate key processes observed in asthma patients. This is achieved by selecting models that mimic the disease’s natural progression, including relevant sensitization routes, doses, and allergens. These models are also instrumental in evaluating potential therapeutic interventions and investigating biomarkers, inflammatory mediators, and distinct asthma endotypes (38).

Bacterial infections have been implicated in asthma exacerbations in human patients. Intracellular bacteria like *Chlamydia pneumoniae* have been associated with fixed airflow limitation (39–41), while *Haemophilus influenzae* has been identified in the sputum of individuals with severe asthma (42). More recently, severe asthma (SA) patients have exhibited neutrophilic airway inflammation alongside infections with *Moraxella catarrhalis* and bacterial species from the *Haemophilus* and *Streptococcus genera* (43). These bacteria can grow intracellularly and produce the second messenger cyclic-di-GMP (c-di-GMP) (44), a potent mucosal adjuvant that promotes Th1-Th17 immune responses while suppressing Th2 responses (45).

Although the impact of Sema3E has been investigated in the acute and chronic HDM models of asthma (14–16, 19, 20, 22, 23, 25), which represent type-2 high asthma in humans, its role in type-2 low or neutrophilic asthma using cyclic-di-GMP remains unexplored. In this study, we investigated Sema3E’s role in a type-2 low asthma model and compared the results with a type-2 high model. Our findings reveal the dichotomous and selective homeostatic roles of Sema3E in asthma models with distinct severities, immune niches, and inflammatory responses.

In this study, the deletion of Sema3E significantly altered inflammatory cell recruitment and cytokine production across different asthma models. *Sema3E KO* mice exhibited enhanced inflammatory cell recruitment, with a more pronounced effect in the type-2 low model, where neutrophils, dendritic cells, and CD4+ T cells were significantly increased compared to WT mice. In contrast, eosinophils were more prevalent in the type-2 high model. Cytokine analysis revealed that *Sema3E KO* mice had elevated pro-inflammatory cytokines, like TNF and type-1 cytokines (IL-17A and IFN-γ) in the type-2 low model and increased IL-4 levels in the type-2 high model, highlighting Sema3E’s role in modulating distinct inflammatory pathways. While total and HDM specific IgG1 and IgE levels remained unchanged, *Sema3E KO* mice exhibited an increase in IgG1-producing B cells in the type-2 low model and IgE-producing B cells in the type-2 high model compared to WT counterpart, suggesting a regulatory role of Sema3E in antibody production. Additionally, our unpublished data suggests that Sema3E plays a critical regulatory role in germinal center responses and IgE production at steady state and post-HDM immunization, independent of ICOS/ICOS-L signaling. These findings emphasize the negative regulatory role of Sema3E in germinal center responses, antibody production, and allergic inflammation.

Inflammation is a central feature of asthma that contributes to its chronicity and severity (46). The differences in immune cell profiles between the type-2 low and type-2 high models underscore the complexity of asthma phenotypes (47). The predominance of neutrophils and type-1 cytokines (IFNγ and IL-17) in the type-2 low model suggests a more aggressive inflammatory response and compromise lung function, as we observed three-fold increase in total lung resistance and five-fold increase in the lung total cell numbers compared to naïve mice, this higher severity is reflecting severe type-2 low phenotype in asthmatic patients (5) In contrast, the eosinophilic response, accompanied by elevated Th2 cytokines (IL-4, IL-5, and IL-13) in the type-2 high model, showed slightly milder response compared to type-2 low model, as we observed two-fold increase in the total lung resistance and lung total immune cells, reflects type-2 high driven inflammation in asthmatics patients with mild to moderate symptoms (5, 47). This distinction suggests that Sema3E plays a differential regulatory role depending on the immunological context, which has implications for targeted therapies.

The Sema3E-plexinD1 axis is known to regulate both neutrophils and eosinophils in allergic asthma (14, 20, 22, 26). Studies indicate that Sema3E-deficient mice exhibit increased neutrophil influx into the lungs in both acute and chronic HDM models of asthma (20, 22). Similarly, in this study, we observed a higher neutrophil count in the lungs of *Sema3E KO* mice in the type-2 low model compared to WT mice. The therapeutic administration of recombinant Sema3E has been shown to reduce neutrophil migration into the airways, decrease CXCL-8 and IL-17 levels *in vivo*, and inhibit CXCL-8/IL-8-mediated human neutrophil migration *in vitro* (20). The elevated IL-17 and CXCL8 levels in *Sema3E KO* mice in the type-2 low model further support its role in neutrophilic inflammation, not only in HDM asthma, as previously reported (20), but also in type-2 low asthma, where IL-17 and CXCL8 are key cytokines in neutrophil recruitment and activation in severe asthma (36).

Beyond neutrophils, Sema3E may inhibit eosinophilic inflammation by blocking lung neovascularization or reducing adhesion molecule expression on eosinophils and eotaxin-1/CCL11 expression on endothelial cells (14). The increased IL-4, IL-5, and IL-13 levels in the type-2 high model and specifically IL-4 increased levels in response to Sema3E deletion compared to WT, suggest that loss of Sema3E creates a microenvironment favorable to eosinophil recruitment, further exacerbating allergic inflammation. While the exact mechanisms by which Sema3E regulates granulocytes remain unclear, it is evident that this axis influences neutrophil and eosinophil migration and inflammatory responses in asthma.

The increased number of DCs and CD4⁺ T cells in the lungs of Sema3E-deficient mice in the type-2 low asthma model, compared to WT mice, highlights the broader impact of this axis on immune regulation beyond granulocytes. DCs are essential for antigen presentation and shaping the adaptive immune response (47). Their accumulation in *Sema3E KO* mice suggests an exaggerated immune activation, which aligns with the observed increase in CD4⁺ T cells. These T cells are key drivers of Th1/Th17 responses, which are characteristic of type-2 low inflammation (47). Thus, the absence of Sema3E appears to enhance DC recruitment and activation, contributing to excessive T cell-driven inflammation.

The elevated levels of IFN-γ and IL-17 in *Sema3E KO* mice correlate with enhanced DC-driven T cell activation. In type-2 low asthma, DCs produce IL-12 and IL-23, which promote Th1 and Th17 differentiation (5, 48). Th1 cells secrete IFN-γ, which enhances macrophage activation and perpetuates airway inflammation, while Th17 cells produce IL-17, a cytokine known to recruit and activate neutrophils in severe asthma (36, 48). The absence of Sema3E likely facilitates increased DC recruitment and activation, amplifying antigen presentation and promoting excessive CD4⁺ T cell responses (22, 23, 25). These findings suggest that Sema3E serves as a key regulator of DC-mediated immune modulation in type-2 low asthma.

In this study, we observed higher mucus production and goblet cell hyperplasia in the type-2 high model compared to the type-2 low model of asthma. Moreover, Sema3E deletion exacerbated goblet cell hyperplasia and mucus production in the type-2 high model, suggesting that Sema3E primarily regulates mucus overproduction selectively based on the allergen regimen. The elevated mucus production and goblet cell hyperplasia in the type-2 high model are likely driven by its characteristic Th2-dominant inflammatory response, which includes high levels of IL-4, IL-5, IL-13 and IL-9 cytokines known to promote goblet cell differentiation and mucus production in airway epithelial cells (5, 36). In contrast, the type-2 low model, which exhibits a more neutrophilic or mixed inflammatory profile, is less dependent on type-2 cytokines-driven goblet cell expansion (5, 36), explaining the comparatively lower mucus production observed.

In this study, Sema3E deficiency significantly influences airway responsiveness and lung function in different asthma models. *Sema3E KO* mice displayed increased AHR and sensitivity to methacholine, especially in the type-2 low asthma model. Notably, airway resistance was higher in WT mice compared to *Sema3E KO* mice in this model, suggesting that the airways in *Sema3E KO* mice may not respond to methacholine increasing doses, indicating enhanced hypersensitivity rather than hyperreactivity in type-2 low model. Although PC20 analysis in type-2 low model between *Sema3E KO* and WT did not reach the statistical significance, there was a reducing trend in *Sema3E KO* mice, suggesting the hypersensitivity in response to Sema3E deletion in the airways. Additionally, *Sema3E KO* mice exhibited significantly greater tissue resistance and elastance in type-2 low model, highlighting more severe lung tissue obstruction and stiffness compared to WT mice. In the type-2 high model, although there was heightened bronchoconstriction and tissue resistance, the significant difference was only in tissue resistance between *Sema3E KO* and WT mice, indicating higher resistance to airflow in *Sema3E KO* mice. Lastly, based on the fold increase in the total lung resistance, the type-2 low model considered more severe compared to type-2 high model. These findings suggest that Sema3E plays a crucial role in modulating airway responsiveness and lung function, with its deficiency exacerbating airway dysfunction more prominently in type-2 low model of asthma compared to type-2 high model, indicating distinct regulatory roles in the structural and functional changes within the lungs across these phenotypes.

The distinction between type-2 low and type-2 high asthma models is crucial. These results indicate that *Sema3E KO* mice exhibited more pronounced bronchoconstriction and impaired lung function in the type-2 low model compared to type-2 high. This finding supports the hypothesis that different immunological contexts can significantly alter the impact of regulatory networks like Sema3E on airway responses (14). This variability underscores the complexity of asthma phenotypes and suggests a need for tailored therapeutic approaches.

The Sema3E-plexinD1 axis plays a crucial regulatory role in airway hyperresponsiveness (AHR) and other key features of asthma (19). In *Sema3E KO* mice challenged with HDM, AHR parameters, including tissue elastance, tissue resistance, and airway resistance, were significantly elevated following methacholine exposure compared to WT controls (22, 23). Sema3E has been shown to inhibit growth factors that drive human airway smooth muscle cell (ASMC) proliferation and migration *in vitro* (24), which may contribute to the heightened AHR observed in *Sema3E KO* mice after HDM exposure (19, 23).

Given the complexity of asthma phenotypes and the regulatory role of Sema3E, Future studies should investigate whether blocking PlexinD1 signaling, Sema3E canonical receptor, *in vivo* mitigates granulocyte recruitment or whether enhancing Sema3E expression selectively in immune cells reduces airway inflammation. Moreover, future research should investigate its interactions with other signaling molecules to identify novel therapeutic strategies for both type-2 low and type-2 high asthma.

Overall, our findings highlight that Sema3E selectively regulates AHR, immune responses, cytokine production, and mucus secretion depending on the immunological niche. This underscores its crucial homeostatic role in modulating both immune and structural changes across distinct asthma phenotypes. Moreover, the fact that Sema3E consistently showed the homeostatic and regulatory role across three different mouse strains (129P2 (22, 25, 26), C57BL/6 (unpublished data), and BALB/c (16, 23)) and four different asthma models (acute (2 weeks) (16, 23), chronic (11 weeks) (22), type 2-low (4 weeks), and type 2-high (4 weeks)) is notable. This consistency across genetic backgrounds and disease phenotypes suggests that Sema3E plays a fundamental role in modulating airway inflammation and remodeling. Furthermore, these findings may provide valuable insights into the heterogeneity of human asthma, as they reflect variations in immune responses, disease severity, and underlying pathophysiological mechanisms observed in patients.

## Acknowledgment

The authors would like to thank Dr. Christine Zhang (Flow Cytometry Core Facility, University of Manitoba) for her help with flow cytometry experiments.

## Authorship

Conception and design: ASG, MM. Acquisition of data and analysis: MM, FS, LS, SB. Interpretation of data: MM, ASG. Draft the article: MM. Critically review the article: MM, ASG Final approval: MM, FS, LS, SB, AH, ASG.

## Abbreviation

Sema3E: Semaphorin-3E HDM House dust mite
AHR: Airway hyperresponsiveness
c-di-GMP: Cyclic dimeric guanosine monophosphate
BALF: Bronchoalveolar fluid
PAS: Periodic acid Schiff
H&E: Hematoxylin and eosin
SA: Severe asthma
ASMC: Human airway smooth muscle cell

